# Role of DEPDC1 in Glioblastoma Malignant Phenotype and Radiosensitivity

**DOI:** 10.1101/2025.05.25.655961

**Authors:** Yang Yu, Hui Deng, Ping Song, Yushu Liu, Qingya Qiu, Mengxian Zhang

## Abstract

Radiotherapy has significantly improved survival outcomes in glioblastoma (GBM) patients, particularly when combined with tumor resection and temozolomide chemotherapy. However, radioresistance remains a major obstacle, limiting further advances in prognosis and potential cure. DEPDC1, a gene implicated in various malignant phenotypes, has not yet been investigated for its role in GBM radioresistance, despite suggestive evidence from previous studies. In this study, analysis of the cancer database and GBM tissue microarrays revealed that DEPDC1 expression is significantly elevated in GBM tissues compared to normal brain tissues. Using GBM cell lines with DEPDC1 knockdown, we observed markedly reduced proliferation and motility. Notably, DEPDC1 downregulation significantly enhanced radiosensitivity, as demonstrated by increased radiation-induced apoptosis, G2/M phase arrest, DNA damage, and impaired DNA repair post-radiation, resulting in lower survival rates in clonogenic assays. Mechanistically, these effects may be mediated through DEPDC1-driven upregulation of NF-κB, supported by tumor sample analysis from our xenograft mouse model. Collectively, our findings suggest that DEPDC1 is not only involved in GBM malignancy but also represents a promising therapeutic target for overcoming radioresistance and developing effective radiosensitizers.

## Introduction

Glioblastoma multiforme (GBM) is the most malignant tumor in the central nervous system, characterized by a poor prognosis (five-year survival rate <10%) [1]. According to the NCCN guidelines, standard therapy for GBM includes surgical resection followed by radiotherapy and temozolomide chemotherapy [2]. However, surgical resection is often limited due to the tumor’s critical location within the brain, and many chemotherapeutic agents are unable to effectively cross the blood-brain barrier [3]. Radiotherapy has been established as an effective modality for GBM treatment. A Surveillance, Epidemiology, and End Results (SEER) analysis of 21,783 patients from 1973 to 2007 showed that radiotherapy alone yielded significantly better survival outcomes than surgical resection alone, with patients receiving postoperative radiotherapy living an average of seven months longer than those who did not receive it [4]. Unfortunately, despite advancements in radiotherapy technology, the prognosis for GBM has not improved significantly, which may be attributed to the acquired radioresistance in GBM [5]. Therefore, elucidating the underlying mechanisms of GBM radioresistance is crucial for developing targeted therapies aimed at overcoming this therapeutic challenge.

DEP domain containing 1 (DEPDC1, also known as DEPDC1A) is a highly conserved mammalian protein encoded by the DEPDC1A gene, initially identified in bladder cancer [6]. DEPDC1 is highly expressed in several malignancies, including bladder, liver, and lung cancers, while its expression is low in most normal tissues [6-8]. Moreover, DEPDC1 has been reported to play a critical role in various cancers. Knockdown of DEPDC1 in bladder cancer and multiple myeloma suppresses cell proliferation and induces apoptosis [6, 9]. In nasopharyngeal carcinoma, DEPDC1 promotes cell proliferation, mediates cell cycle progression, and supports cell motility [10]. In prostate cancer, DEPDC1 may interact with SEDC23B to regulate cell viability, invasion, and apoptosis [11]. In breast cancer immunotherapy research, DEPDC1 has been proposed as a potential immunogenic epitope [12]. In lung adenocarcinoma, DEPDC1 promotes proliferation and inhibits apoptosis via NF-κB signaling pathway [13]. In glioma, DEPDC1 knockdown similarly suppressed tumor proliferation and induced apoptosis [14]. Collectively, these findings suggest that DEPDC1 may serve as a key oncogenic factor in GBM.

Previous studies showed that DEPDC1 downregulates A20 transcription through forming a transcriptional repressor complex with ZNF224 [13, 15, 16]. Since A20 inhibits the phosphorylation of IκB-α, A20 downregulation leads to NF-κB activation that contributes to cell proliferation, DNA repair and resistance to apoptosis [17]. Given that these NF-κB–mediated effects contribute to radioresistance in GBM [18, 19], DEPDC1 may potentially enhance GBM radioresistance via the NF-κB signaling pathway. Additionally, DEPDC1 has been reported to regulate cell cycle progression, particularly at the G2/M phase [10, 20, 21]. As cells in G2/M phase are extremely sensitive to radiation (RT) [22], targeting DEDPC1, a G2/M phase regulator, may represent a promising strategy to enhance the efficacy of radiotherapy.

Therefore, our study focuses on investigating the role of DEPDC1 in GBM, particularly its impact on radiosensitivity, which may direct the development of novel radiosensitizers for GBM radiotherapy.

## Materials and Methods

### Cell Culture

Human GBM cell lines U251 and U87 were obtained from the American Type Culture Collection (ATCC). Cells were maintained and cultured in Dulbecco’s Modified Eagle Medium (DMEM, Hyclone) supplemented with 10% fetal bovine serum (Gibco) at 37°C in a humidified atmosphere containing 5% CO2.

### Bioinformatics Analysis

The transcript expression of DEPDC1 across various cancer types was analyzed using the validated datasets from the Human Protein Atlas (HPA), which are derived from The Cancer Genome Atlas (TCGA) and include data for 10 cancer types. To assess DEPDC1 expression in normal brain tissue, low-grade gliomas, and GBM, expression profiles were obtained from the Gene Expression Omnibus (GEO) database. The dataset GSE4290 was selected for this analysis.

### Tissue Microarray and Immunohistochemical (IHC) Staining

A GBM tissue microarray was obtained from Xi’an Alenabio Technology Co. Ltd., consisting of 70 GBM tissue samples and 10 normal or adjacent brain tissue samples. All tissues were initially fixed in formalin and embedded in paraffin. DEPDC1 expression was assessed by immunohistochemical (IHC) staining. The paraffin-embedded tissue microarray was baked at 60°C for 30 minutes, then deparaffinized and rehydrated. Antigen retrieval was performed using 10 mM citrate buffer (pH 6.0), followed by blocking with 10% fetal bovine serum in TBS. Slides were incubated with a rabbit anti-DEPDC1 antibody (Sigma; 1:50 dilution), and subsequently with an anti-rabbit secondary antibody (Dako Cytomation). Staining was visualized using DAB and counterstained with hematoxylin. Images were captured and quantified using an expression level scoring system. Scores were determined by multiplying the staining intensity score (0 = negative, 1 = weak, 2 = moderate, 3 = strong) by the percentage of positive staining area (0 = 0%, 1 = 1–25%, 2 = 26–50%, 3 = 51–75%, 4 = >75%).

### Lentiviral Construction and Packaging

The shRNA sequence targeting DEPDC1 (Patent CN111269910B) and a scrambled control sequence were cloned into plasmids. These plasmids were selectively amplified in E. coli and subsequently inserted into the GV248 lentiviral vector (GeneChem), which contains an enhanced green fluorescent protein (EGFP) reporter gene, an internal ribosome entry site (IRES), and a puromycin resistance gene. Lentiviral particles were produced by transfecting 293T cells and stored at –80°C for later use. The sequences of the DEPDC1-targeting shRNA (shDEPDC1) and control shRNA (shCtrl) were as follows: shDEPDC1: TATCCAGTAAGGCTATCAT; shCtrl: TTCTCCGAACGTGTCACGT

### Transfection and Isolation of Stable Cell Clones

To knockdown DEPDC1, cells were transfected with lentiviruses carrying either shCtrl or shDEPDC1 using Polybrene (Beyotime). Following transfection, cells were allowed to recover for 72 hours, after which the culture medium was replaced with selection medium containing 2.5 μg/mL puromycin for 48 hours. EGFP-positive cells were then observed under a fluorescence microscope. If transfection efficiency did not exceed 90%, the selection process was repeated. DEPDC1 knockdown was validated by RT-PCR and Western blotting.

### Real-Time Reverse Transcription–Polymerase Chain Reaction (RT-PCR)

Total RNA was extracted from cells or tissues using DNAiso Reagent (Clontech) according to the manufacturer’s instructions. RNA was reverse-transcribed and amplified using the SYBR Green RT-PCR Kit (Quanti-Tect). Relative gene expression levels were calculated using the ΔCt method, normalizing the Ct values of target genes to that of the reference gene GAPDH. The primers used were as follows: DEPDC1 forward: 5’-ATGCGTATGATTTCCCGAATGAG-3’; DEPDC1 reverse: 5’-CACAGCATAACACACATCGAGAA-3’; GAPDH forward: 5’- TGACTTCAACAGCGACACCCA-3’; GAPDH reverse: 5’-CACCCTGTTGCTGTAGCCAAA-3’; RELA forward: 5’-ATGTGGAGATCATTGAGCAGC-3’; RELA Reverse : 5’- CCTGGTCCTGTGTAGCCATT-3’.

### Western Blotting (WB)

Cells or tissues were harvested, homogenized, and lysed with ice-cold RIPA buffer containing 1% PMSF (Beyotime). Protein concentrations were determined using the BCA protein assay. Equal amounts of protein from each sample were separated by SDS-PAGE and transferred onto polyvinylidene difluoride (PVDF) membranes. The membranes were blocked with 5% powdered milk for 1 hour and incubated overnight at 4°C with primary antibodies against DEPDC1 (Mouse, Abcam), p65 (Rabbit, Proteintech), GAPDH (Mouse, Abcam), or β-actin (Rabbit, Abcam) at a dilution of 1:1000. After washing the membranes three times with TBST, they were incubated with corresponding secondary antibodies (Anti-Rabbit or Anti-Mouse, Boster) at a dilution of 1:5000 for 1 hour. Immunoblots were visualized using the Bio-RAD Blot Imaging System.

### Celigo Cell Counting

Cells were seeded into 96-well plates at a density of 2000 cells per well and cultured in a 37°C incubator. The cell number in each well was assessed daily using the Celigo Imaging Cytometer (Revvity). Proliferative curves were generated based on the daily measurements.

### Clonogenic Assay

Cells were counted and seeded into 6-well plates at the following densities: 500 cells per well for 0 Gy, 500 cells per well for 2 Gy, and 1000 cells per well for 4 Gy. The plates were irradiated with doses ranging from 0 to 4 Gy using RS 2000 series Biological Irradiator (Rad Source Technologies). After irradiation, cells were incubated at 37°C for 10 to 14 days. Colonies were stained with crystal violet, and those containing at least 50 cells were counted by microscopic inspection.

### Wound Healing Assay

Cells were seeded into 6-well plates and cultured until a confluent monolayer formed. A linear scratch was created across the cell layer using a 20□μL pipette tip, and the wells were washed three times with PBS to remove detached cells. The medium was then replaced with serum-free (starvation) medium. Initial images of the wound area were captured, and corresponding images of the same region were taken after 16 hours. Cell migration was evaluated by measuring the distance migrated into the wound area using microscopic analysis.

### Flow Cytometry Analysis

Apoptosis was assessed using the Annexin V-PE/7-AAD apoptosis detection kit (BD Biosciences) according to the manufacturer’s instructions. Cells were seeded in 6-well plates and treated with or without 8 Gy radiation. After 72 hours, 1×10□ cells from each group were harvested, resuspended in 100□μL of 1× Binding Buffer, and stained with Annexin

V-PE and 7-AAD for 15 minutes at room temperature. Samples were analyzed by flow cytometry within 1 hour, and data were processed using FlowJo software. For cell cycle analysis, 1×10□ cells from each group were collected 72 hours post-irradiation (8 Gy), fixed in 70% ice-cold ethanol, and stored at 4°C. After washing with PBS, cells were treated with RNase A for 30 minutes and stained with propidium iodide (PI). DNA content was analyzed by flow cytometry, and the proportions of cells in G1, S, and G2/M phases were determined using ModFit software.

### Immunofluorescence (IF)

Cells were seeded on coverslips in 24-well plates and incubated for 24 hours. They were then treated with or without 6 Gy radiation, either 1 hour or 24 hours before fixation. Cells were fixed in 4% ice-cold paraformaldehyde for 15 minutes and permeabilized with 0.5% Triton X-100 for 10 minutes. After blocking with 5% BSA for 30 minutes, cells were incubated overnight at 4°C with an anti-γ-H2AX antibody (Rabbit, Cell Signaling Technology; 1:400). The following day, cells were incubated in the dark for 1 hour with a Cy3-labeled goat anti-rabbit secondary antibody (Boster; 1:500). Nuclei were counterstained with DAPI, and coverslips were mounted using Antifade Mounting Medium (Boster). Images were acquired using a confocal fluorescence microscope.

### Comet Assay

Cells were harvested 24 hours after irradiation with 6 Gy. Cells were then suspended in 130□μL of 0.8% low melting-point agarose and layered onto fully frosted slides pre-coated with 1% normal melting-point agarose. Slides were then incubated in alkaline lysis buffer (4□mol/L NaCl, 500□mmol/L Na_2_-EDTA, 500□mmol/L Tris-HCl, 1% sodium N-lauroyl sarcosinate, 1% Triton X-100, and 10% DMSO; pH 10) at 4°C overnight in the dark. Next, slides were immersed in electrophoresis buffer (1□mmol/L Na_2_-EDTA and 300□mmol/L NaOH) for 30 minutes to allow DNA unwinding, then subjected to electrophoresis at 25□V and 300□mA for 30 minutes. Slides were subsequently neutralized in 0.4□mol/L Tris buffer (pH 7.5) for 5 minutes and fixed in methanol for 10 minutes. DNA was stained with SuperRed (Biosharp), and comet images were captured using a fluorescence microscope. DNA damage was quantified using the Olive Tail Moment (OTM), calculated as: OTM = Tail Moment Length × Tail % DNA.

### Animal Study

Twenty female BALB/c nude mice (4 weeks old; Shanghai Lingchang Biotechnology Co., Ltd., China) were randomly assigned to two groups: negative control (NC) and knockdown (KD). A total of 5×10□ U87 cells (shCtrl or shDEPDC1) were subcutaneously injected into the right forelimb of each mouse. After 19 days, tumor volume was measured every 3 days using calipers and calculated using the formula: Volume = (length × width^2^ × π) / 6. On day 33, all mice were sacrificed, and tumors were excised and weighed. Tumor tissues were collected for further analysis by RT-PCR and WB.

### Statistical Analysis

Statistical analyses were performed using GraphPad Prism or Microsoft Excel. Data are presented as mean ± standard deviation (SD) unless otherwise indicated. Comparisons between groups were conducted using Student’s t-test or two-way ANOVA. A *p* < 0.05 was considered statistically significant.

## Results

### DEPDC1 is highly expressed in GBM but not in normal brain tissue

Previous studies have shown that DEPDC1 exerts cancer-promoting effects in various malignancies [6-14], including lung, liver, and breast cancers. Herein, we evaluated DEPDC1 transcript expression across multiple cancer types using data from HPA (Human Protein Atlas proteinatlas.org) [23]. Among the ten cancer types included in the HPA dataset, GBM exhibited the highest average DEPDC1 expression (**Fig. 1A**). Notably, DEPDC1 expression in GBM was significantly higher than in lung, liver, and breast cancers (*p* < 0.0001), where its tumor-promoting role has already been established, suggesting an even more prominent involvement of DEPDC1 in GBM. To assess DEPDC1 expression across different brain tissue types, mRNA levels were analyzed using expression profiles from the GEO dataset GSE4290, which included 23 normal brain tissue samples, 76 grade II–III glioma samples, and 81 GBM samples [24, 25]. DEPDC1 mRNA was significantly upregulated in GBM tissues compared to both normal brain tissues (fold change = 5.64, *p* < 0.0001) and low-grade gliomas (fold change = 2.38, *p* < 0.0001). In comparison, low-grade gliomas exhibited only a modest increase in DEPDC1 expression relative to normal tissues (fold change = 2.37, *p* = 0.029), underscoring the association between DEPDC1 overexpression and GBM malignancy (**Fig. 1B**).

**Figure 1:**
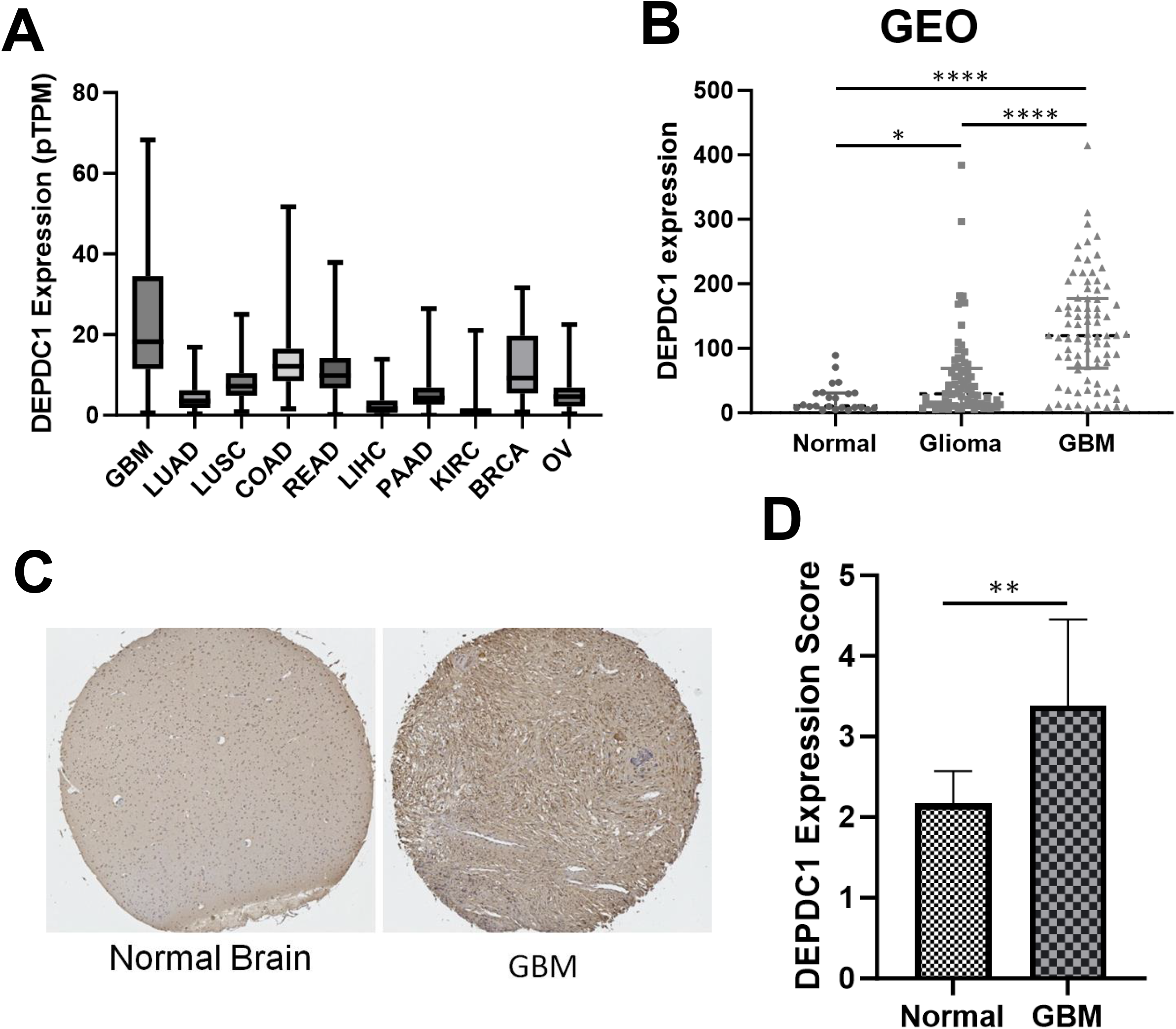
DEPDC1 expression is elevated in GBM. **(A)** Boxplot showing DEPDC1 mRNA expression across various cancer types based on data from HPA. GBM shows the highest average expression; Student’s t-test vs. all other cancers, *p*□<□0.0001. LUAD, lung adenocarcinoma; LUSC, lung squamous cell carcinoma; COAD, colon adenocarcinoma; READ, rectum adenocarcinoma; LIHC, liver hepatocellular carcinoma; PAAD, pancreatic adenocarcinoma; KIRC, kidney renal clear cell carcinoma; BRCA, breast invasive carcinoma; OV, ovarian serous cystadenocarcinoma. **(B)** DEPDC1 mRNA expression levels in normal brain tissues, Glioma (Grade II-III), and GBM samples from the GEO database GSE4290. **(C)** Representative IHC staining images of DEPDC1 in normal brain tissue and GBM samples from a tissue microarray. **(D)** Quantification of DEPDC1 IHC expression scores in normal brain and GBM tissues. Statistical significance was determined using Student’s t-test. \**p* < 0.05, \*\**p* < 0.01, \*\*\**p* < 0.001, \*\*\*\**p* < 0.0001.

To validate these findings at the protein level, DEPDC1 expression was evaluated by IHC using a tissue microarray comprising GBM samples (n = 70) and normal brain tissue samples (n = 10). As shown in **Fig. 1C–D**, the average DEPDC1 expression score in GBM tissues was approximately 30% higher than in normal tissues (*p* = 0.007), confirming that DEPDC1 is also upregulated at the protein level in GBM.

### Generation of stable DEPDC1 knockdown GBM cell lines

To investigate the role of DEPDC1 in GBM, two GBM cell lines (U87 and U251) were selected for the establishment of stable isogenic cell lines with DEPDC1 knockdown. These cell lines were specifically chosen due to their high malignancy, elevated DEPDC1 expression, and their proven suitability for transfection, making them ideal candidates for studying the functional effects of DEPDC1 depletion in GBM [26, 27]. The parental cell lines were transduced with lentiviral vectors expressing DEPDC1-targeted shRNA or a scrambled control shRNA, both linked to a puromycin resistance gene and an EGFP reporter. Following puromycin selection, more than 90% of the cells expressed EGFP (**Fig. 2A**), confirming efficient transfection. RT-PCR and WB analyses further demonstrated that DEPDC1 was significantly downregulated at both the mRNA and protein levels in both cell lines (**Fig. 2B-D**), validating the stable establishment of DEPDC1 knockdown isogenic cell lines.

**Figure 2:**
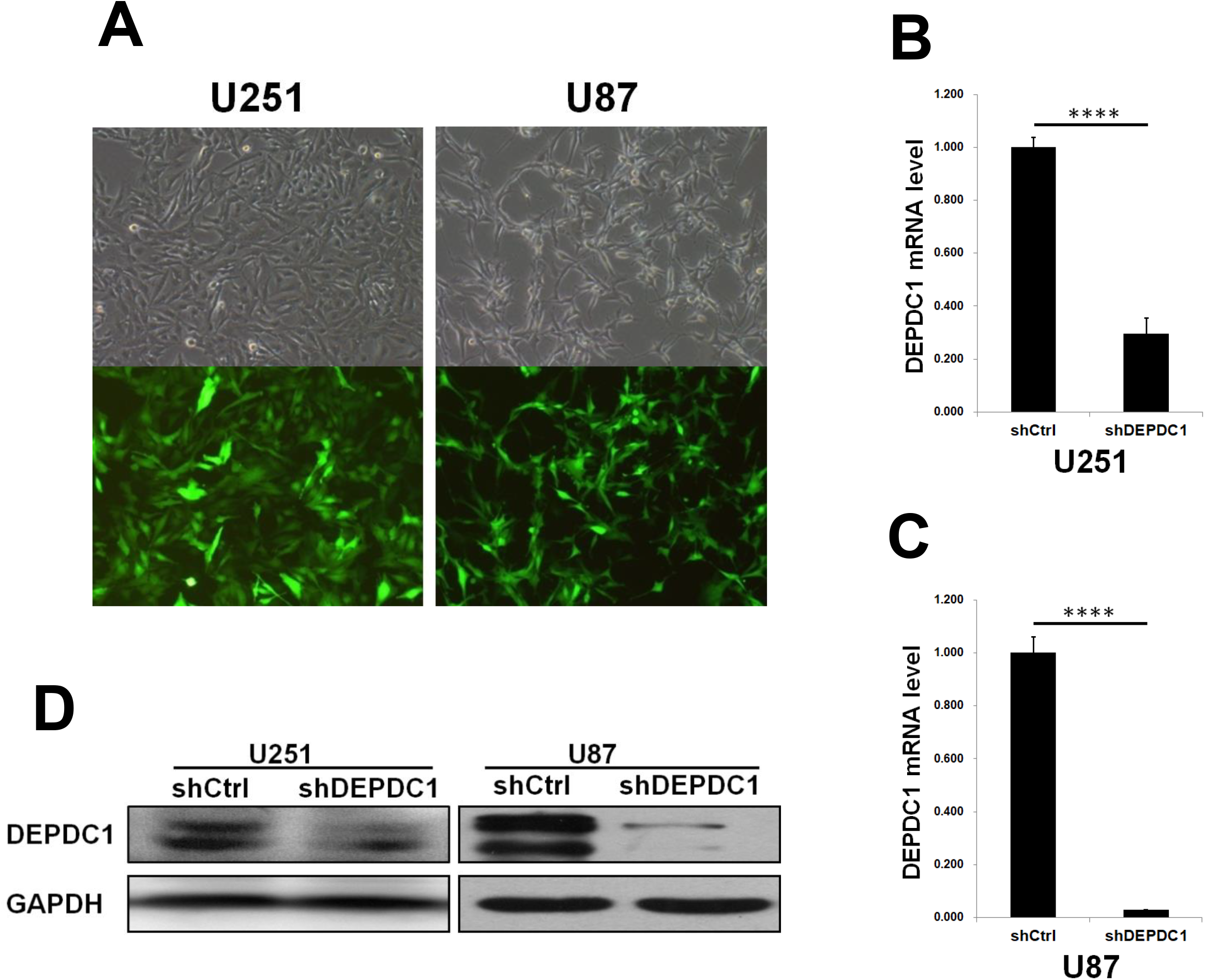
Validation of transfection in U87 and U251 isogenic cell lines. **(A)** Fluorescence microscopy images of GBM cells transfected with lentiviruses carrying the EGFP gene. **(B-C)** DEPDC1 mRNA expression levels in U251 **(B)** and U87 **(C)** measured by RT-PCR. **(D)** DEPDC1 protein expression detected by WB. Statistical significance was determined using Student’s t-test. \*\*\*\**p* < 0.0001.

### DEPDC1 knockdown inhibits cell proliferation and motility in GBM

As demonstrated in previous studies [6, 9-11, 13, 14], DEPDC1 promotes cell proliferation in various types of cancers, suggesting its oncogenic role. To assess its functional relevance in GBM, we monitored cell proliferation using the Celigo Imaging Cytometer in established DEPDC1 knockdown cell lines. Quantitative analysis of proliferation curves showed a marked reduction in cell growth upon DEPDC1 knockdown in both U87 and U251 cells. Two-way ANOVA revealed a significant divergence between DEPDC1-silenced and control groups in both cell lines (*p* < 0.0001), indicating that DEPDC1 promotes GBM cell proliferation (**Fig. 3A-B**).

**Figure 3:**
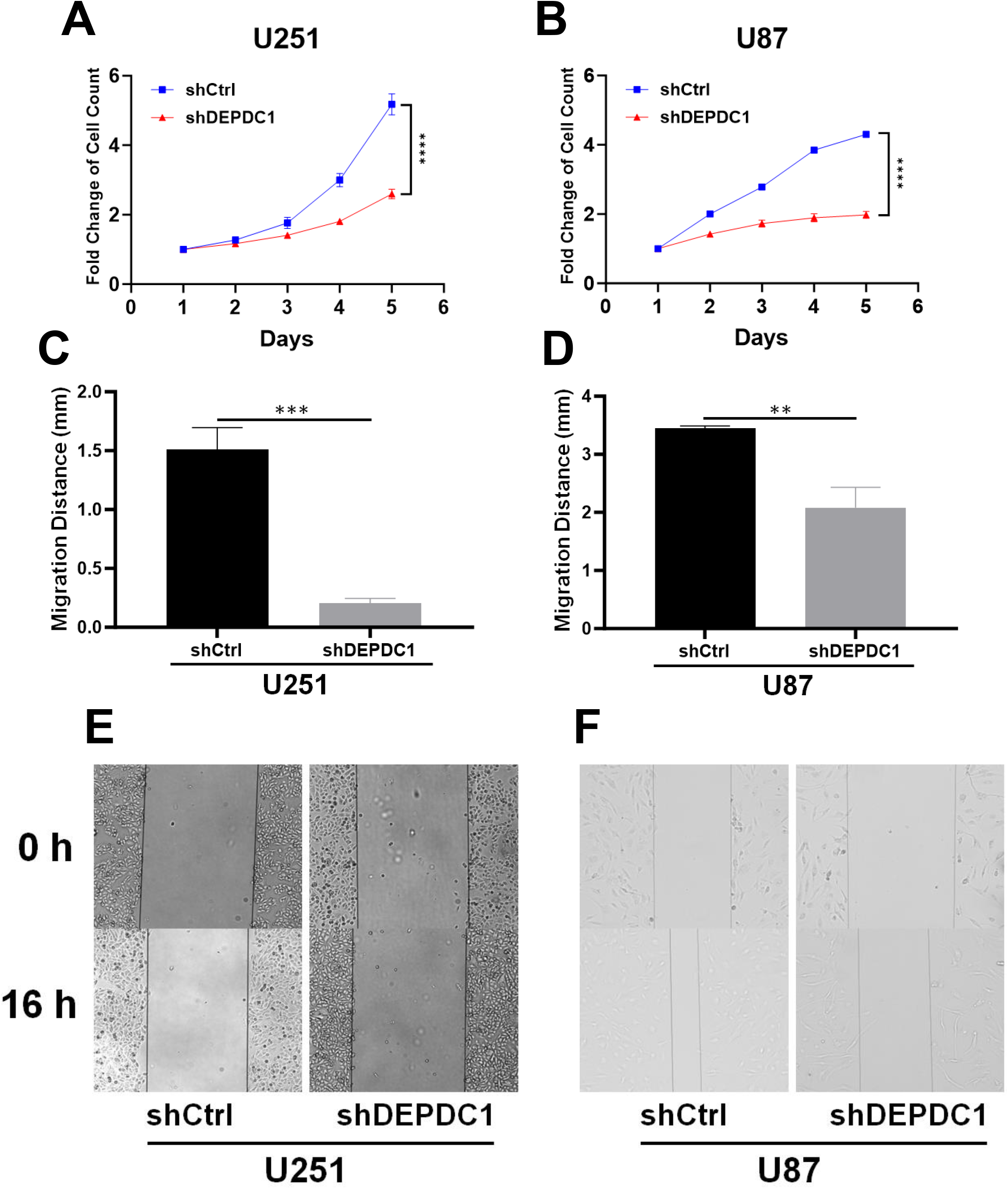
DEPDC1 promotes GBM cell proliferation and motility. **(A-B)** Cell proliferation of U251 **(A)** and U87 **(B)** cells measured by Celigo cell counting over 5 days. Curves were compared using two-way ANOVA. **(C-D)** Quantification of cell migration for U251 **(C)** and U87 **(D)** cells. Migration distance was measured at 16 h. **(E-F)** Representative images of wound healing assays at 0 h (top) and 16 h (bottom) in U251 **(E)** and U87 **(F)** cells. Statistical significance was determined using Student’s t-test **(C–D)**. \*\**p* < 0.01, \*\*\**p* < 0.001, \*\*\*\**p* < 0.0001.

Epithelial-mesenchymal transition (EMT) is a key mechanism that contributes to radioresistance in cancer cells [28], often associated with increased cell motility [29]. Thus, we further determined the role of DEPDC1 in GBM cell motility. A wound healing assay was performed on confluent monolayers of U87 and U251 cells with or without DEPDC1 knockdown. To specifically assess migration independent of proliferation, the assay was conducted in serum-free (starvation) medium. Sixteen hours after scratch induction, DEPDC1-knockdown cells exhibited significantly reduced wound closure compared to controls in both U87 and U251 (*p* < 0.05), indicating impaired motility (**Fig. 3C-F**). These findings suggest that DEPDC1 facilitates both proliferative and migratory capacities in GBM cells, potentially contributing to tumor aggressiveness.

### DEPDC1 knockdown enhances the suppressive effects of radiotherapy on GBM survival

To assess the role of DEPDC1 in modulating GBM cell sensitivity to radiation, U87 and U251 isogenic cell lines (shCtrl vs. shDEPDC1) were exposed to increasing doses of X-ray irradiation (0 Gy, 2 Gy, and 4 Gy). Clonogenic assays, the gold standard for evaluating radiation-induced reproductive cell death [30], were performed to determine post-RT survival and estimate the potential for tumor recurrence. As shown in Figures **4A–F**, both DEPDC1 knockdown and RT significantly reduced clonogenic survival in U87 and U251 cells (*p* < 0.0001 across all doses). Though RT alone inhibited colony formation in a dose-dependent manner in both U87 and U251 cells, DEPDC1 knockdown further amplified this suppressive effect (**Fig. 4C-D**). At 2 Gy, colony formation inhibition was enhanced more substantially in DEPDC1-deficient cells than in controls (U251: 64.3% vs. 45.3%; U87: 45.0% vs. 34.4%). A similar trend was observed at 4 Gy (U251: 95.6% vs. 88.5%; U87: 75.3% vs. 68.2%). From another perspective, while DEPDC1 knockdown alone significantly attenuated colony-forming capacity in both cell lines (*p* < 0.01), its inhibitory effect was further strengthened in the presence of RT (**Fig. 4E-F**). In U251 cells, DEPDC1 knockdown significantly reduced clonogenic survival by 36.9% without RT, which increased to 58.9% and 76.0% at 2 Gy and 4 Gy, respectively. Similarly, the suppression increased from 29.9% (0 Gy) to 41.3% (2 Gy) and 45.5% (4 Gy) in U87 cells. Collectively, these results indicate that DEPDC1 knockdown acts synergistically with RT to inhibit GBM cell survival, highlighting its potential role in modulating the cellular response to radiation.

**Figure 4:**
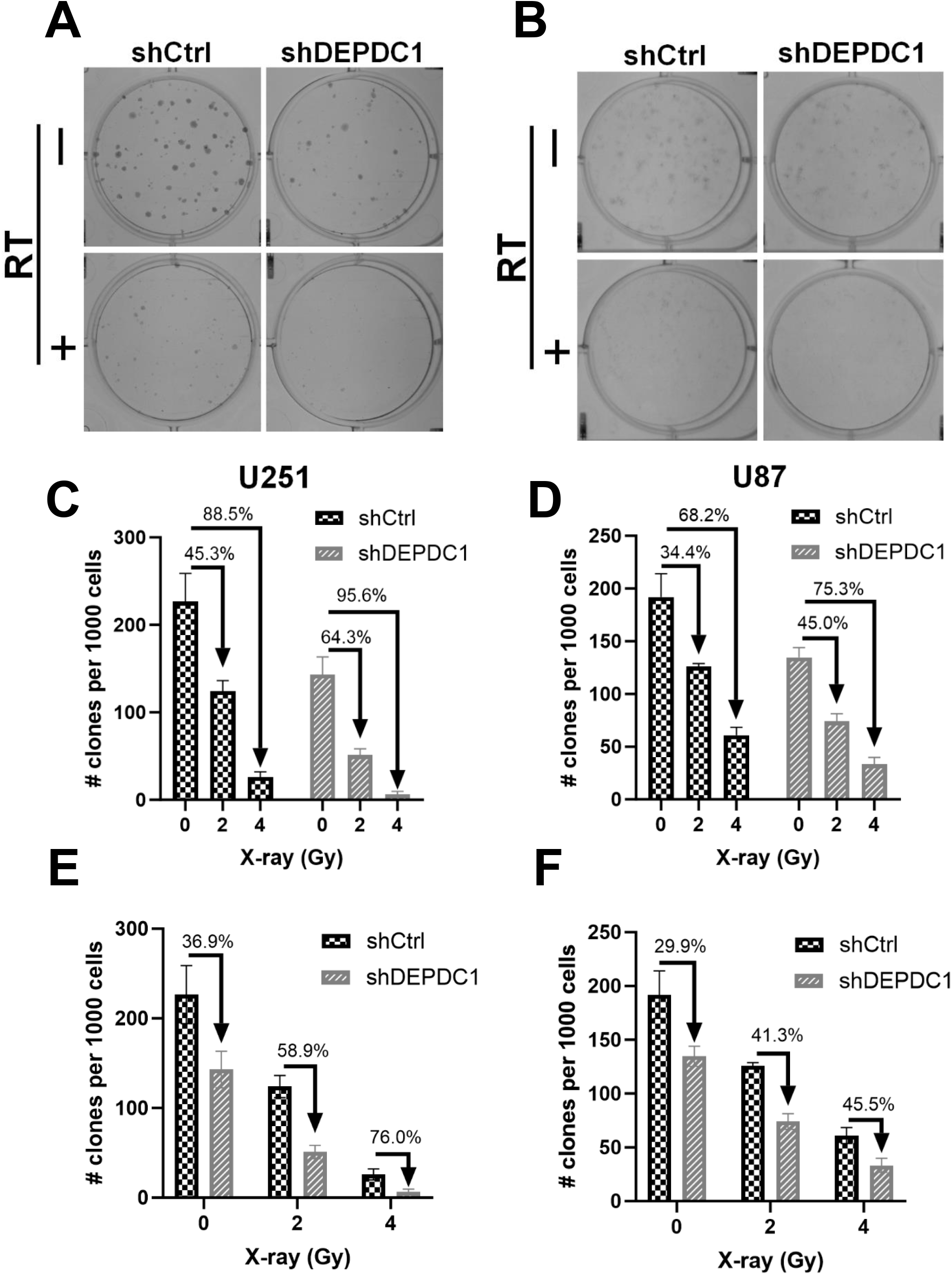
DEPDC1 knockdown reduces GBM cell survival following RT. **(A-B)** Representative images of colony formation in U251 **(A)** and U87 **(B)** cells with or without DEPDC1 knockdown following RT. **(C-F)** Quantification of clonogenic survival in U251 **(C, E)** and U87 **(D, F)** cells exposed to 0, 2, or 4 Gy X-ray. Colony numbers were normalized to 1000 seeded cells. Two-way ANOVA showed that both DEPDC1 knockdown and radiation dose significantly reduced colony formation (*p* < 0.0001).

### DEPDC1 knockdown enhances radiation-induced apoptosis and G2/M phase arrest in GBM cells

Building upon the observed synergistic effects between DEPDC1 knockdown and RT on clonogenic survival, we next investigated the underlying cellular mechanisms contributing to this enhanced radiosensitivity. Radiation-induced apoptosis is a key contributor to radiotherapeutic efficacy in cancer cells [31]. To assess whether DEPDC1 knockdown augments apoptotic responses to radiation, U87 and U251 cells transduced with either shCtrl or shDEPDC1 were exposed to 8 Gy of X-ray irradiation. Apoptosis levels were quantified using Annexin V-PE/7- AAD double staining followed by flow cytometry, both prior to and 72 hours post-RT. In U251 cells (**Fig. 5A, C**), RT alone caused a modest, non-significant increase in early apoptosis at 72 h post-RT. In contrast, combined DEPDC1 knockdown and RT significantly elevated early apoptotic cell levels compared to both untreated and irradiated shCtrl cells (*p* < 0.01). In U87 cells (**Fig. 5B, D**), RT alone significantly increased the proportion of 7-AAD_+_ cells, indicating late apoptosis or necrosis (*p* < 0.05), but did not significantly affect early apoptosis. Notably, the combination of DEPDC1 knockdown and RT not only significantly increased the 7-AAD_+_ population (*p* < 0.01), but also led to a significant rise in early apoptotic cells compared to RT alone (*p* < 0.05). DEPDC1 knockdown alone had no significant apoptotic effect in either cell line under non-irradiated conditions. Collectively, these findings suggest that DEPDC1 knockdown sensitizes GBM cells to RT-induced apoptosis, particularly by enhancing early apoptotic responses.

**Figure 5.**
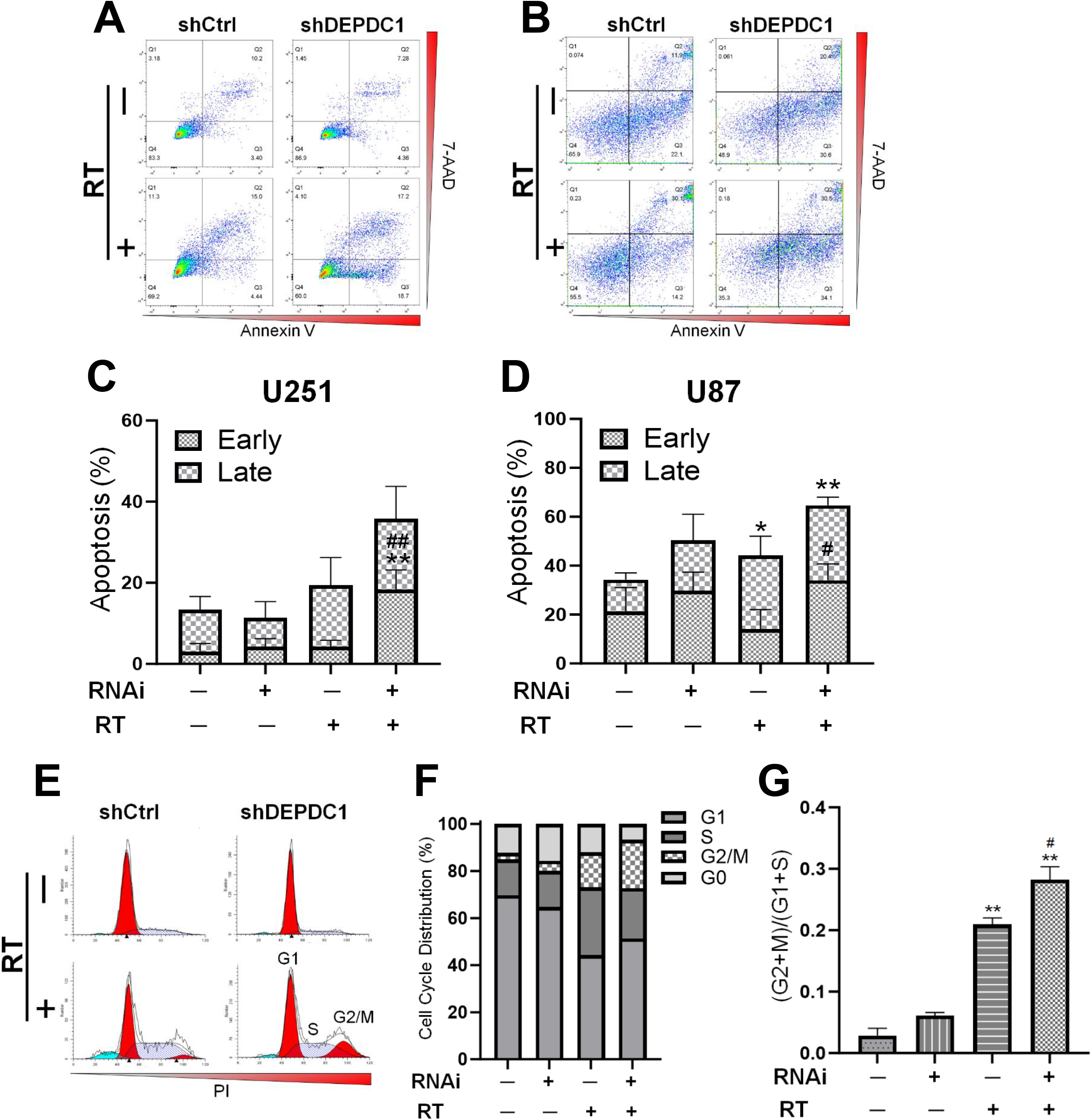
DEPDC1 knockdown enhances RT-induced apoptosis and G2/M arrest. **(A-B)** Representative flow cytometry plots of Annexin V/7-AAD staining in U251 **(A)** and U87 **(B)** cells. Q2 (Annexin-V^+^,7-AAD^+^), late apoptosis or necrosis; Q3 (Annexin-V^+^,7-AAD^-^), early apoptosis; Q4 (Annexin-V^-^,7-AAD^-^), viable cells. **(C-D)** Quantification of the apoptotic cells in U251 **(C)** and U87 **(D)**. Early, early apoptosis; Late, late apoptosis or necrosis. **(E)** Representative histograms of cell cycle distribution in U251 cells. **(F)** Quantification of cells in G1, S, and G2/M phases. **(G)** Ratios of G2/M to G1+S phases were compared across groups. RNAi indicates DEPDC1 knockdown. Statistical significance was determined using Student’s t-test. \**p* < 0.05, \*\**p* < 0.01 compared to shCtrl without RT. ^#^*p* < 0.05, ^##^*p* < 0.01 compared to shCtrl with RT.

In addition to apoptosis, cell cycle arrest, particularly at the G2/M checkpoint, is another well-documented mechanism of RT-induced growth inhibition [31]. To determine the role of DEPDC1 in this context, U251 cells expressing shCtrl or shDEPDC1 were subjected to 8 Gy irradiation, followed by cell cycle analysis via 7-AAD staining and flow cytometry. As depicted in **Fig. 5E–G**, While RT alone significantly induced G2/M phase arrest (*p* < 0.01), DEPDC1 knockdown led to a significant increase in the proportion of cells arrested in the G2/M phase following RT, compared to shCtrl without RT (*p* < 0.01) or with RT (*p* < 0.05). No significant differences in cell cycle distribution were observed between groups under non-irradiated conditions. These results suggest that DEPDC1 depletion facilitates radiation-induced G2/M arrest, potentially contributing to reduced proliferation and enhanced apoptosis.

### DEPDC1 knockdown enhances radiation-induced DNA damage and inhibits DNA repair in GBM cells

Given the observed enhancement of RT-induced apoptosis and cell cycle arrest by DEPDC1 knockdown, we next investigated whether these effects were associated with altered DNA damage responses. Since RT primarily exerts its cytotoxic effects through the induction of DNA double-strand breaks, and cancer cells often rely on DNA repair mechanisms to counteract this genotoxic stress [32, 33]. We therefore examined whether DEPDC1 influences DNA damage levels and repair efficiency following RT. To assess DNA damage, a comet assay was performed in U251 isogenic cells (shCtrl vs. shDEPDC1) following 6 Gy irradiation. The results indicated both RT and DEPDC1 knockdown individually increased Olive tail moments (*p* < 0.001 and *p* < 0.05, respectively), indicative of DNA strand breaks. Notably, cells with DEPDC1 knockdown exhibited significantly greater Olive tail moments following RT compared to shCtrl cells with or without RT (*p* < 0.0001 for both) (**Fig. 6A-B**), suggesting that DEPDC1 knockdown exacerbates RT-induced DNA damage.

**Figure 6.**
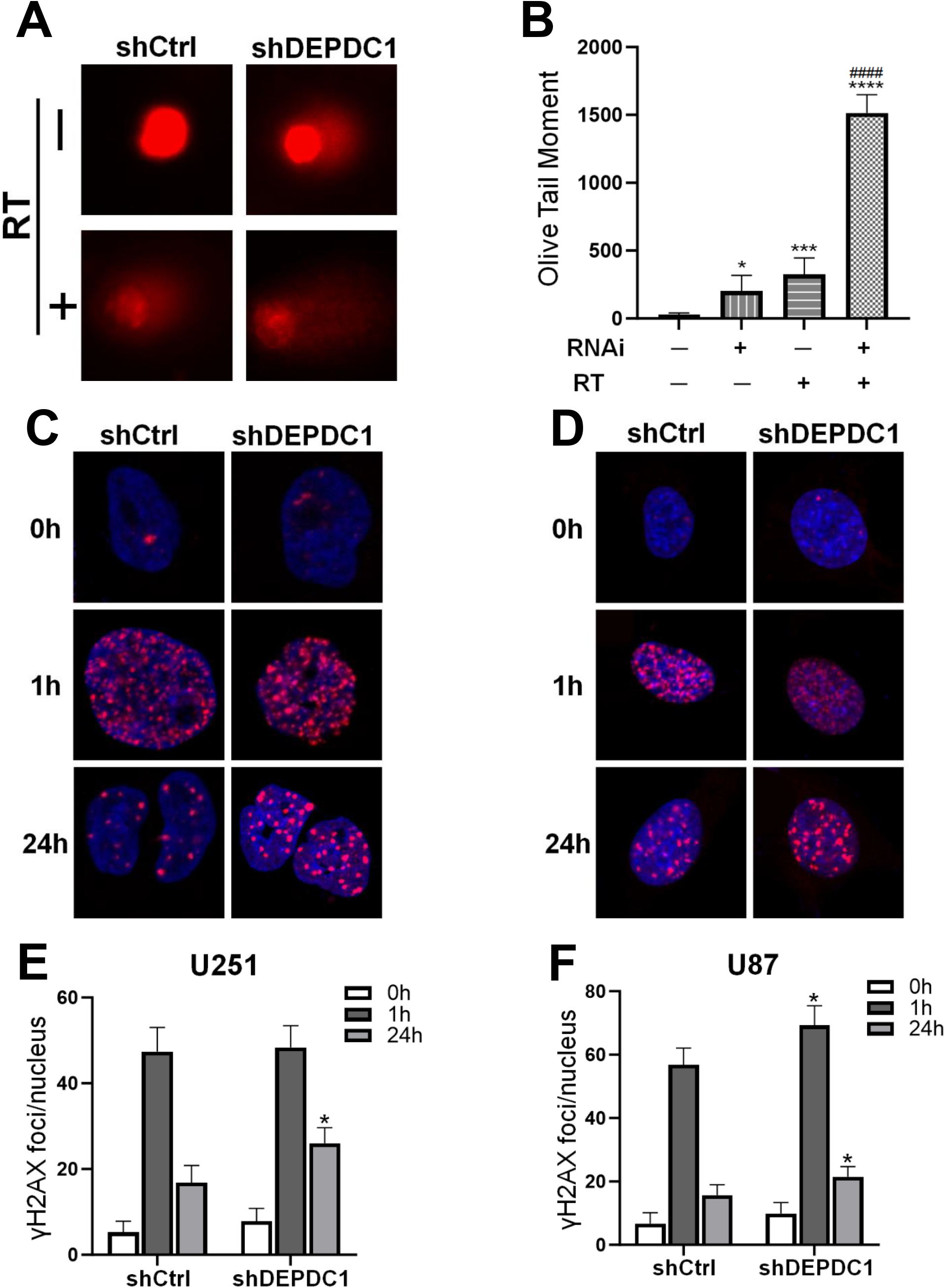
DEPDC1 knockdown enhances RT-induced DNA damage and impairs repair. **(A)** Representative images of alkaline comet assays in each group. **(B)** Quantification of OTM scores. RNAi indicates DEPDC1 knockdown. Statistical significance was determined using Student’s t-test. \**p* < 0.05, \*\*\**p* < 0.001, \*\*\*\**p* < 0.0001 vs. shCtrl without RT; ^####^*p* < 0.0001 vs. shCtrl with RT. **(C-D)** Representative IF images of γ-H2AX foci (red) and nuclei (DAPI, blue) in U251 **(C)** and U87 **(D)** cells. **(E-F)** Quantification of γ-H2AX foci per nucleus in U251 **(E)** and U87 **(F)** at indicated time points. Student’s t-test was used to compare shCtrl and shDEPDC1 groups at the same post-RT time point. \**p* < 0.05 vs. shCtrl at the corresponding time point.

To further evaluate the dynamics of DNA damage and repair, we assessed the formation and resolution of γ-H2AX foci, widely used markers of DNA double-strand breaks [34], via IF in U87 and U251 cells. In both cell lines, exposure to 6 Gy of RT led to a sharp increase in the number of γ-H2AX foci per nucleus at 1 hour post-RT, confirming efficient induction of DNA damage. By 24 hours post-RT, γ-H2AX foci levels decreased substantially in cells, reflecting ongoing DNA repair processes. However, DEPDC1-knockdown cells persisted a significantly higher number of γ-H2AX foci at 24 hours post-RT compared to vector controls (*p* < 0.05 in both cell lines) (**Fig. 6C–F**), indicating impaired DNA damage resolution. Together, these findings demonstrate that DEPDC1 knockdown not only intensifies RT-induced DNA damage but also hinders the subsequent repair, which may underlie the increased apoptosis and G2/M arrest observed earlier.

### DEPDC1 knockdown inhibits tumor growth and downregulates NF-κB expression *in vivo*

To investigate the role of DEPDC1 in glioblastoma progression *in vivo*, twenty nude mice were randomized into either the control (NC) or DEPDC1 knockdown (KD) group followed by 5×10^6^ U87 isogenic cells (shCtrl or shDEPDC1, respectively) subcutaneous injection. Tumor growth was monitored every three days starting from day 19 post-inoculation using caliper measurements. As shown in **Fig. 7B**, tumors in the KD group exhibited significantly slower growth compared to the NC group. At the endpoint (day 33), both the average tumor volume and weight were markedly reduced in the KD group (**Fig. 7A,C**). These results suppose that DEPDC1 knockdown suppresses GBM growth *in vivo*.

**Figure 7.**
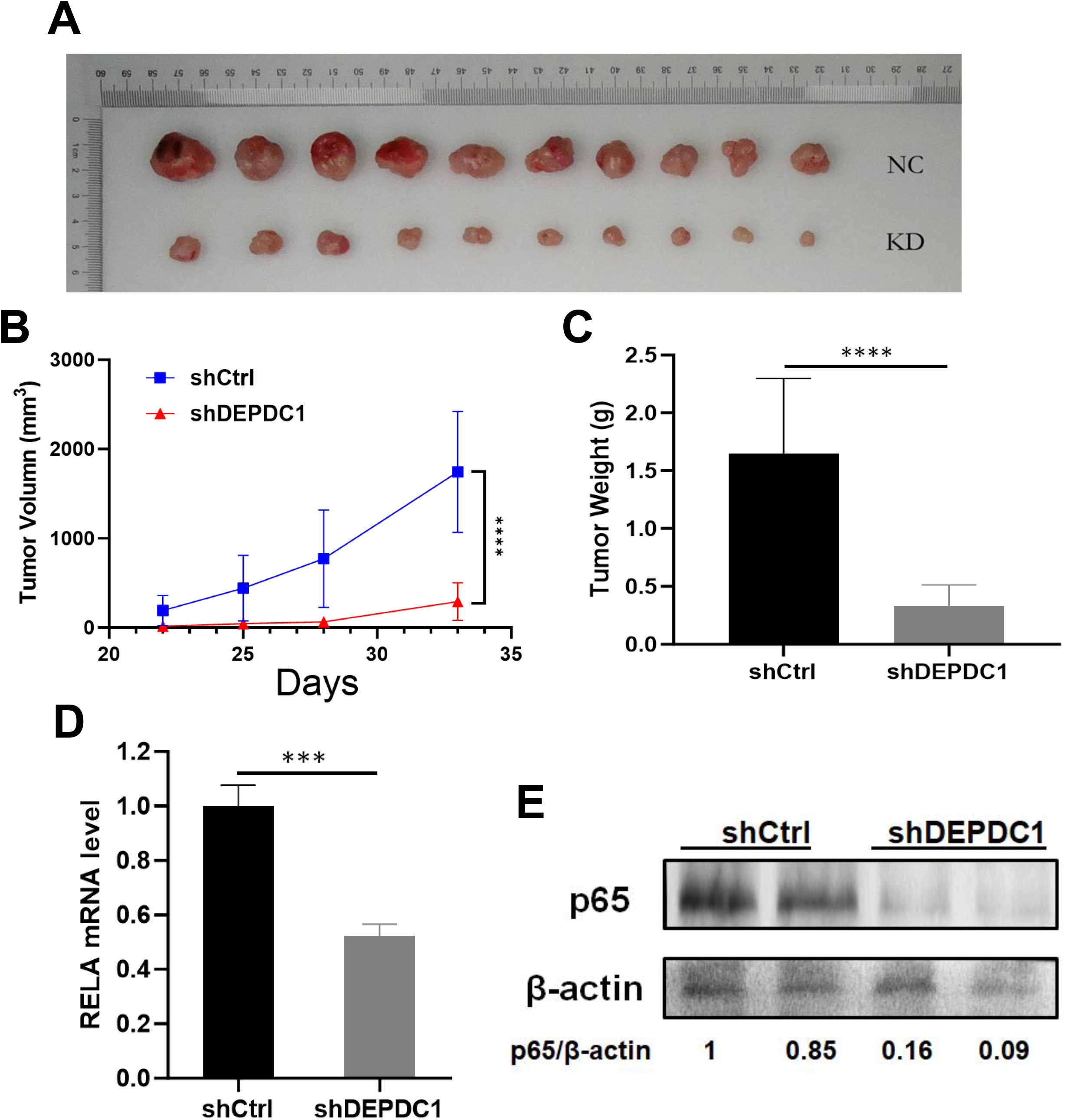
Effects of DEPDC1 knockdown on GBM tumor growth in a nude mouse xenograft model. **(A)** Photographs of xenografts harvested 33 days after subcutaneous injection. **(B)** Tumor volumes were measured every three days. Two-way ANOVA was used to compare growth curves. \*\*\*\**p* < 0.0001. **(C)** Average tumor weights at endpoint. **(D)** RELA mRNA expression in xenografts was assessed by RT-PCR. Student’s t-test was used for comparisons in **(C)** and **(D)**. \*\*\**p* < 0.001, \*\*\*\**p* < 0.0001. **(E)** Protein levels of NF-κB p65 subunit in xenografts were analyzed by WB. Band intensities were quantified, and p65/β-actin ratios were calculated for each sample.

Given that NF-κB may serve as a marker of radioresistance, and has been reported to be upregulated by DEPDC1 in several previous studies [17-19, 35], we assessed the expression of RELA (encoding the p65 subunit of NF-κB [36]) at the mRNA and protein levels in tumor tissues. RT-PCR and WB revealed reduced RELA mRNA and p65 protein expression in the DEPDC1 knockdown group compared to controls (**Fig. 7D-E**). These results suggest that DEPDC1 may promote GBM progression and radioresistance, at least in part, through upregulation of the NF- κB (p65) pathway.

## Discussion

DEPDC1 has been identified as a promising therapeutic target in various types of cancer, contributing to key oncogenic processes such as tumorigenesis, tumor progression, and cell cycle dysregulation [6, 9-14]. Mechanistically, DEPDC1 is thought to promote tumor growth mainly through the DEPDC1–A20–NF-κB signaling axis [13, 15, 16]. NF-κB is a pivotal transcription factor involved in the regulation of various processes in glioblastoma (GBM), including the maintenance of cancer stem-like cells, promotion of mesenchymal identity, enhancement of invasive potential, and resistance to radiotherapy [37]. As an upstream regulator of NF-κB, DEPDC1 may potentially contribute to radioresistance in GBM. To our knowledge, this study provides the first systematic evidence supporting a role for DEPDC1 in cancer radioresistance.

In line with previous findings [14] and supported by our data, DEPDC1 knockdown alone was able to induce a modest level of apoptosis in GBM cells, potentially due to impaired repair of DNA replication errors [38]. However, this effect was relatively limited and may not be sufficient for therapeutic efficacy when used as monotherapy. In contrast, when DEPDC1 knockdown was combined with radiotherapy, we observed a markedly enhanced anti-tumor effect. Given that radiotherapy remains a cornerstone of GBM treatment, our findings suggest that DEPDC1 inhibition may serve as a valuable radiosensitizing strategy rather than a stand-alone therapeutic approach.

The observed enhancement of RT-induced G2/M phase arrest following DEPDC1 knockdown is likely multifactorial. One explanation is that DEPDC1 silencing impairs DNA damage repair, possibly via suppression of NF-κB signaling. However, additional mechanisms may also contribute. DEPDC1 has been implicated in centrosome integrity and spindle organization during mitosis [21], and its depletion may disrupt proper chromosome segregation, predisposing cells to G2/M arrest. Additionally, previous studies indicate that DEPDC1 may regulate the expression of cyclins CCNB1 and CCNB2, which control the G2 to M phase transition [20, 39]. Interestingly, DEPDC1 knockdown has been associated with upregulation of CCNB1 and CCNB2, potentially pushing more cells into the division cycle and increasing their susceptibility to G2/M arrest after RT [40]. These hypotheses warrant further investigation to elucidate the precise mechanisms by which DEPDC1 regulates the cell cycle.

Beyond its effects on tumor proliferation, DEPDC1 also appears to influence GBM cell motility. One possible explanation is that DEPDC1 enhances NF-κB activity, which plays unique roles in neural tissue, including the regulation of synaptic plasticity and dendritic spine development [41]. Although direct mechanisms remain unclear, studies on CCR7 and SLC8A2 suggest that NF-κB signaling may modulate GBM cell migration. Specifically, CCR7 has been shown to promote TGF-β1-mediated NF-κB activation associated with increased migration, while SLC8A2 suppresses migration by inhibiting NF-κB signaling [42, 43]. Similarly, DEPDC1 might influence cell motility, at least in part, through NF-κB–related pathways. Furthermore, DEPDC1 may also affect motility independently of NF-κB, potentially through its involvement in cytoskeletal regulation, as indicated by its reported interaction with anti-tubulin agents [44].

A noteworthy concern lies in the complex and context-dependent role of A20, a key regulator within the DEPDC1–A20–NF-κB axis. While A20 is generally regarded as a tumor suppressor, it has paradoxically been shown to support glioma stem cell maintenance [45]. This raises the possibility that DEPDC1 knockdown may restore A20 expression, which could support the survival of glioma stem cells while inhibiting growth in non-stem tumor cells. Given that glioma stem cells are known to drive recurrence and confer resistance to radiotherapy [46], caution must be exercised when considering DEPDC1 as a therapeutic target. Further studies are needed to clarify the divergent roles of the DEPDC1–A20 axis in GBM stem versus non-stem cell populations.

## Conclusion

DEPDC1 is highly expressed in GBM and promotes tumor growth and migration. DEPDC1 knockdown enhances radiosensitivity by increasing apoptosis, inducing G2/M arrest, and impairing DNA repair. In vivo, DEPDC1 knockdown reduces tumor growth and NF-κB expression, suggesting its potential as a radiosensitizer target in GBM treatment.

## Authors Contributions

Conceptualization: Y.Y. and M.X.Z. Original draft writing, preparation of figures and tables: Y.Y. and M.X.Z. Data acquisition and analysis: Y.Y., H.D., P.S., Y.S.L., and Q.Y.Q. Revising and editing: Y.Y. and M.X.Z. Supervision: M.X.Z. All authors have read and agreed to the published version of the manuscript.

## Acknowledgements

We acknowledge funding support for this study from National Natural Science Foundation of China (NSFC). Grant number: 81772680.

